# Inhibition of Sphingosine-1-Phosphate signaling in pancreatic ductal adenocarcinoma leads to downregulation of growth and invasion

**DOI:** 10.1101/2020.09.29.318964

**Authors:** Brenna A. Rheinheimer, Alex Cardenas, Luis Camacho, Evan S. Ong, Tun Jie, Ronald L. Heimark

**Affiliations:** Department of Surgery, The University of Arizona, Tucson, AZ, USA; Division of General Surgery, Western Surgical Group, Reno, NV, USA; Department of Medicine, The University of Arizona, Tucson, AZ, USA; Division of General Surgery, Swedish Medical Group, Seattle, WA, USA; The University of Arizona Cancer Center, The University of Arizona, Tucson, AZ, USA

## Abstract

**Background:** Deregulated phosphorylation of sphingosine by the sphingosine kinases and signaling through the EDG family of receptors enhances growth and survival in many cell types. Therefore, we sought to elucidate the effect of alterations in the ceramide/sphingosine/S1P rheostat on driving human pancreatic ductal adenocarcinoma towards a malignant phenotype.

**Methods:** Pancreatic cancer cell lines were treated with exogenous S1P, FTY720, and siRNA to Sphk1. Migration was evaluated by wound healing assays, cell growth by MTT assays, and invasion by tumorsphere assays. Expression of S1PR1, S1PR3, Sphk1, and Sphk2 were measured by quantitative PCR, western blot, and immunohistochemistry.

**Results:** S1PR1, S1PR3, and Sphk2 were overexpressed in all pancreatic cancer cell lines. Sphk1 translocated from the cytoplasm to the nucleus in cells located at the leading edge of cell clusters. Exogenous S1P increased cell migration while treatment with FTY720 and Sphk1 siRNA decreased cell growth and invasion.

**Conclusions:** Our results suggest that increased S1PR1 expression may be an early event in pancreatic cancer pathogenesis. Additionally, altered Sphk1 localization may provide a mechanism through which pancreatic ductal adenocarcinoma cells at the leading edge invade into the surrounding matrix. Finally, inhibition of sphingosine-1-phosphate signaling may provide a novel therapeutic target for patients with metastatic disease.

## Introduction

Pancreatic ductal adenocarcinoma (PDAC) is the fourth-leading cause of cancer-related death in the United States with a five-year survival rate of 8%^1^. Survival is limited due to the lack of diagnostic markers and late development of clinical symptoms leading to approximately 80% of PDACs being metastatic at the time of diagnosis^2^. Currently, surgery is the only treatment option that renders any long-term survival; however, the five-year survival rate of patients who undergo surgical resection remains a dismal 15-20%^3^. Therefore, it is imperative to further elucidate the genetic landscape underlying PDAC progression and metastasis. Discovery of additional genes contributing to this process would provide advances in the complexity of PDAC invasion and may ultimately lead to the stratification of patients based on the risk of metastatic capabilities and provide novel treatment modalities to patients with metastatic disease.

Sphingolipids are lipid signaling molecules that are associated with cell membranes in eukaryotic cells. Sphingosine-1-phosphate (S1P) is produced by the sphingosine kinases Sphk1 and Sphk2, which catalyze the phosphorylation of sphingosine to S1P. At homeostasis, S1P levels are endogenously low and regulated in a spatial-temporal manner through its production and degradation. Sphk1 localization is mainly cytoplasmic; however, its activation requires translocation to the plasma membrane through phosphorylation on Ser225 by ERK1/2^4^. Sphk2 is located predominantly in the nucleus or on the endoplasmic reticulum^5,6^, but can be exported into the cytosol though phosphorylation by protein kinase D^7^. S1P functions as the natural ligand for the EDG family of G protein-coupled receptors (*S1PR1-5*)^8^. The receptors for S1P are ubiquitously expressed and couple to numerous G proteins leading to a wide range of downstream signaling that controls many diverse biological processes including enhanced growth and survival^9,10^, increased expression of pro-apoptotic proteins and downregulation of anti-apoptotic proteins^11,12^, and inhibition of cytochrome c release^13^. Similar to inositol polyphosphate regulation of chromatin remodeling complexes, S1P regulates transcription by inhibiting histone deacetylase activity of HDAC1 and HDAC2^14^. S1P also promotes cell migration through activation of S1PR1 and S1PR3^15^. Alternatively, neutralization of S1P slowed the progression of lung, breast, melanoma, and ovarian cancers in xenograft mice^16^. Alternatively, increased expression of Sphk1 has been demonstrated to regulate anti-tumor immunity in metastatic melanoma patients^17^. Sphk1 has also been shown to be elevated in solid tumors and protects against apoptosis by inhibiting the intrinsic death pathway through activation of NF-κB^18^. Therefore, the sphingosine-1-phosphate signaling pathway is a promising target to study in pancreatic ductal adenocarcinoma.

The ceramide/sphingosine/S1P rheostat postulates that agents that regulate the interconversion of ceramide/sphingosine/S1P direct the cell towards either apoptosis or survival depending on the relative position of the rheostat. Research has shown that gemcitabine-resistant PDAC cells have a lower ceramide/S1P ratio and increased Sphk1 activity^19^. Exogenous addition of S1P into PDAC cells increased proliferation and migration through the Src signaling pathway^20^. Finally, treatment with FTY720, an S1PR1 antagonist, led to an inhibition of growth, increased apoptosis, and cell cycle arrest in human and mouse PDAC cells^21^. While the sphingosine-1-phosphate signaling pathway has been briefly studied in pancreatic ductal adenocarcinoma, previous research has only looked at the presence of S1P and activity of Sphk1. Therefore, we sought to elucidate the effect of alterations in the ceramide/sphingosine/S1P rheostat on the cellular localization and activity of Sphk1 and Sphk2 and the resulting influence on driving human pancreatic ductal adenocarcinoma towards a more malignant phenotype.

## Materials and Methods

### Cell culture

All pancreatic ductal adenocarcinoma cell lines were obtained from ATCC. BxPC-3, HPAF-II, and Su86.86 were maintained in RPMI 1640 supplemented with 10% heat-inactivated fetal bovine serum and penicillin/streptomycin. PANC-1 were maintained in DMEM supplemented with 10% heat-inactivated fetal bovine serum and penicillin/streptomycin. Capan-1 were maintained in McCoy’s 5A supplemented with 10% heat-inactivated fetal bovine serum and penicillin/streptomycin. Immortalized human pancreatic ductal epithelial cells (HPDE) were obtained from the Ming-Sound Tsao lab at the University of Toronto, Toronto, Ontario, Canada and maintained in keratinocyte serum-free media supplemented with 0.05 mg/ml of bovine pituitary extract and 0.02 µg/ml EGF recombinant human protein. All cells were maintained in a 37°C, 5% CO_2_ atmosphere with constant humidity.

### Nucleic acid isolation and quantitative PCR

Total RNA was extracted using Trizol and quantified by absorption measurements at 260 nm. 1 μg of total RNA was reverse transcribed using 1 μg/ml random hexamer primers and SuperScript II reverse transcriptase. Primers to *S1PR1, S1PR3, SPHK1, SPHK2*, and *GAPDH* were designed using the Roche Universal Probe Library assay design center. Quantitative PCR (qPCR) was performed using SYBR Green Master Mix on an ABI Prism 7500 Sequence Detection System. Differences in expression between cancer cell lines and HPDE were determined using the comparative Ct method described in the ABI user manual relative to *GAPDH* for *S1PR1, S1PR3, SPHK1*, and *SPHK2*. Primer sequences are available upon request.

### Western blot

Monolayers of cultured cells were collected and lysed in 2X SDS sample buffer. Protein concentrations were measured using the BCA method. Nuclear and cytoplasmic fractionation was performed using the Pierce NE-PER fractionation kit per the manufacturer’s instructions. 60 μg of total protein was loaded per well on a 10% polyacrylamide gel for separation and transferred to nitrocellulose membranes by electrophoresis overnight on ice. Membranes were blocked by incubation in 5% nonfat dry milk in TBST for one hour at room temperature. Primary antibodies to S1PR1, Sphk1, and Sphk2 were diluted in either 5% nonfat dry milk in TBST or 5% bovine serum albumin in TBST as directed by the manufacturer. Blots were incubated overnight at 4°C with the primary antibody at dilutions recommended by the manufacturer. Peroxidase-conjugated secondary antibodies were used for signal amplification at a dilution of 1:2000 in 5% nonfat dry milk in TBST for one hour at room temperature. Protein bands were identified by chemiluminescence exposed on X-OMAT AR film. Images of western blots were captured using Metamorph Version 3.0.

### Immunofluorescence

BxPC-3 were grown on glass coverslips to 50% confluence then fixed with 4% para-formaldehyde in PBS for 20 minutes at room temperature. Cells were then extracted on ice with CSK buffer (100 mM NaCl, 300 mM sucrose, 3 mM MgCl_2_, 10 mM PIPES, pH 6.8) for two minutes and blocked (1% bovine serum albumin, 2% normal goat serum) for 30 minutes at room temperature. Primary antibodies were added at dilutions recommended by the manufacturer at 4°C overnight followed by incubation with fluorescent-conjugated secondary antibodies. Cells were counterstained with DAPI for visualization before mounting.

### Wound healing assay

Bx-PC3 were cultured in 35 mm dishes until confluent. Upon confluence, the bottom of the culture dish was scraped once with a 1000 μL pipette tip to create a linear area devoid of cells. One dish was taken immediately for fixation. Remaining dishes were then washed with PBS and supplemented with fresh media containing vehicle control, 100 nM exogenous S1P, or 1000 nM exogenous S1P. After 24 hours, the remaining dishes were washed in PBS and fixed for 30 minutes at room temperature with 4% para-formaldehyde in PBS. After fixation, the dishes were imaged at 5X magnification using confocal microscopy. Images were analyzed by ImageJ to determine the relative area between cells. Results were calculated as a percentage of the area re-colonized by cells compared to the dish taken immediately for fixation.

### MTT assay

PANC-1 cells were plated at a density of 7500 cells per well in a 96-well plate and allowed to attach overnight. 24 hours after plating, cells were supplemented with fresh media containing vehicle control, 0 nM FTY720, 500 nM FTY720, 1000 nM FTY720, 2000 nM FTY720, or 10000 nM FTY720. 24 hours after dosing, 20 μl of prepared MTT was added to each well and cells were incubated at 37°C for 4 hours. After incubation, the solution was aspirated and 150 μl of MTT solvent (4 mM HCl, ipecal in isopropanol) was added to each well. Absorbance was read on an automated plate reader at a wavelength of 562 nm.

### Tumorsphere culture

Cultured PANC-1 cells were trypsinized and re-suspended in complete growth medium containing 0.2% (w/v) of carboxymethylcellulose to a final concentration of 4000 cells/ml. 100 μl of cells were then seeded into non-adherent 96-well plates at a density of 400 cells per well. Plated cells were allowed to settle overnight to form spheroid cultures. Spheroid cultures were then harvested, settled in a 15 ml conical tube, suspended in 200 μl of cold Matrigel, and distributed among four wells of 8 chamber incubation plates. The Matrigel solidified for 30 minutes at 37°C then cells were supplemented with fresh media containing vehicle control, 2 μM FTY720, 5 μM FTY720, and 10 μM FTY720. 24 hours after plating in Matrigel, 5 separate fields were taken on a Zeiss AxioVert 200 microscope at 10X and cells were supplemented with fresh media with/without FTY720. 48 hours after plating in Matrigel, cells were imaged again. The number of projections per spheroid was counted 24 hours and 48 hours after treatment with FTY720.

### Invasion assay

siRNA to Sphk1 was purchased from Santa Cruz Biotechnology and transfected into PANC-1 cells at a final concentration of 50 nmol/L using Oligofectamine transfection reagent. Cells were incubated for 4 hours in OptiMEM serum free media then supplemented with complete growth media. 24 hours after transfection, cells were harvested for further analysis. Matrigel-coated Boyden chambers were rehydrated and 2.5×10^4^ transfected cells were plated in serum free media in the top chamber while the bottom chamber contained complete growth media. 24 hours after plating, the chambers were stained with 50 μl of 0.5% crystal violet in 20% methanol for 1 minute, washed in distilled water, swabbed on the inside to remove noninvasive cells, and allowed to dry overnight. Images of 5 separate fields were acquired using a 40X objective on a Zeiss AxioVert 200 microscope. The number of invading cells per image was counted and data is presented as the mean number of invading cells.

### Immunohistochemistry

Deidentified, formalin-fixed, paraffin-embedded archival tissue blocks were obtained from patients that had not received any preoperative chemotherapy or radiation therapy. All patients provided written consent and all phases of the research were approved by the Internal Review Board of the University of Arizona. A total of 48 different samples were used: 2 adjacent normal, 12 pancreatitis, 11 intraductal papillary mucinous neoplasm (IPMN), and 23 PDAC. Tissue sections were deparaffinized with Xylene and rehydrated. Antigen retrieval was carried out using an Antigen Decloaker. Slides were blocked for one hour at room temperature with 3% normal donkey serum + 0.01% Triton X-100. Primary antibody in 1% normal donkey serum + 0.01% Triton X-100 was added in a 1:25 dilution and slides were incubated overnight at 4°C in a humidified chamber. Biotinylated secondary antibody was added and incubated for 30 minutes at room temperature followed by incubation with Vectastain Elite avidin biotin complex for an additional 30 minutes at room temperature. Slides were developed with 3,3-diaminobenzidine (DAB) and counterstained with hematoxylin. The immunohistochemical stained slides were scanned at 40X magnification on a Zeiss microscope. The number of ductal or tumor cells exhibiting staining was scored as follows based on the number of cells exhibiting staining and the average intensity of staining in those cells. For percentage, the slides were scored as follows: 1 for 0-40% of cells, 2 for 40-70% of cells, 3 for > 70% of cells. For intensity, the scores were 1 (none to light staining), 2 (moderate staining), and 3 (strong staining). A composite score was determined by multiplying the two scores. In addition, cellular localization was determined for SPHK1 and SPHK2.

## Results

### S1PR1, S1PR3, and Sphk2 are overexpressed in pancreatic ductal adenocarcinoma cell lines

S1P signaling through S1PR1 and S1PR3 favors the activation of pro-growth and anti-apoptotic pathways. Therefore, we sought to determine *S1PR1* and *S1PR3* expression in our panel of pancreatic cancer cell lines. BxPC-3, Su.86.86, and PANC-1 showed 2.5-to-4-fold overexpression of *S1PR1* (Figure 1a) while Su.86.86 showed a 7-fold increase and PANC-1 showed a 44-fold increase in *S1PR3* (Figure 1b). All PDAC cell lines showed expression of S1PR1 while HPDE showed no expression (Figure 1c) suggesting that pro-growth signaling in pancreatic ductal adenocarcinoma may occur via S1P signaling through S1PR1. Sphk1 has been shown to increase resistance of PDAC cells to gemcitabine^18^. Therefore, we measured *Sphk1* and *Sphk2* levels in our panel of PDAC cell lines. *Sphk1* was not highly expressed in any of our PDAC cells (Figure 1d); however, *Sphk2* expression was increased 2-to-6.5-fold in all of our PDAC cells (Figure 1e) suggesting that signaling through the sphingosine kinases may play a role in the progression of pancreatic ductal adenocarcinoma.

**Figure 1.**
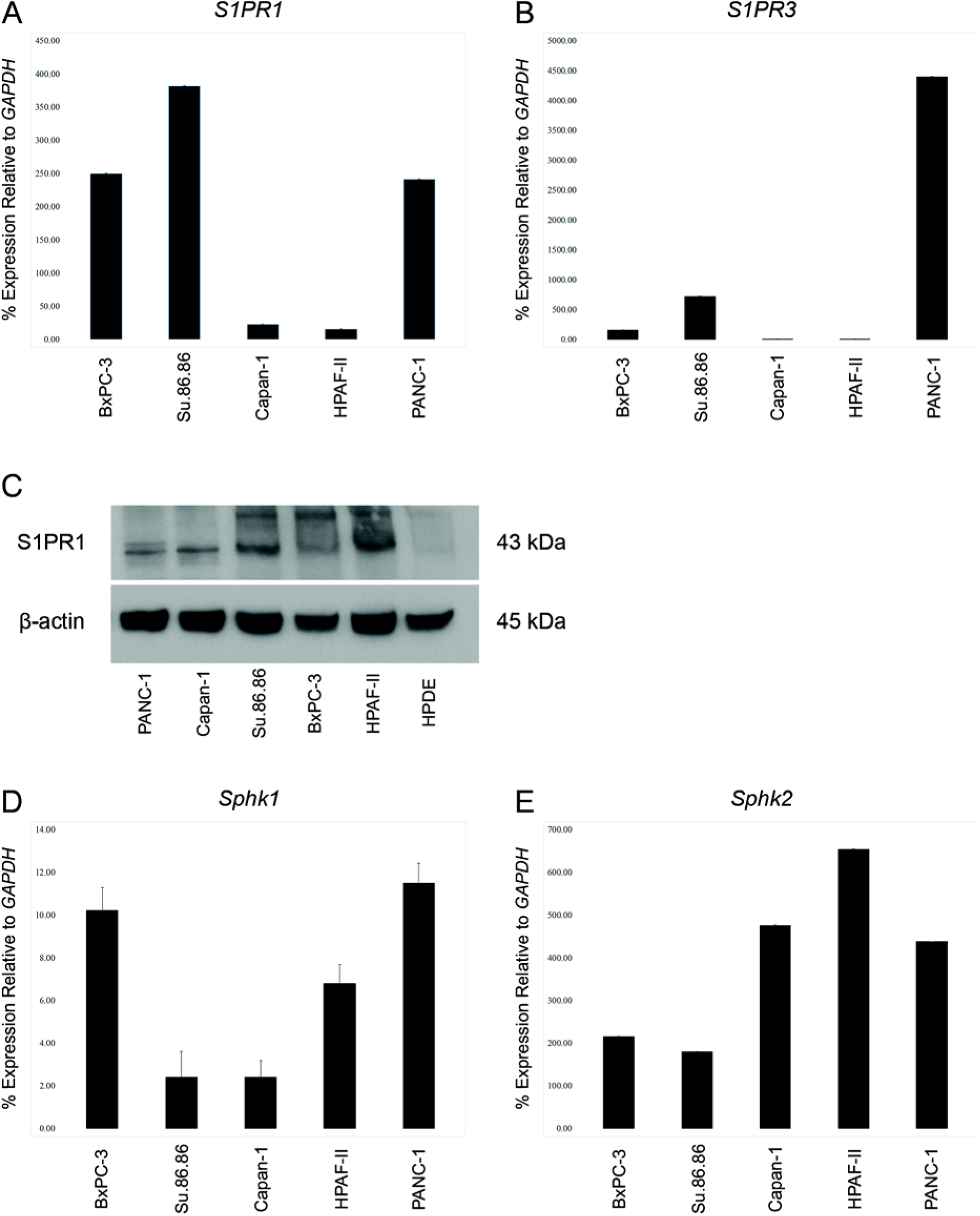
S1PR1, S1PR3, and Sphk2 are overexpressed in pancreatic ductal adenocarcinoma cell lines. (a) qPCR for S1PR1. (b) qPCR for S1PR3. (c) Western blot analysis for S1PR1 protein expression. B-actin was used as a loading control. (d) qPCR for Sphk1. (e) qPCR for Sphk2.

### Sphk1 and Sphk2 localization is altered in pancreatic ductal adenocarcinoma

In other cell types, cytoplasmic localization of Sphk1 and nuclear localization of Sphk2 is necessary for the overall action of Sphk1 to be anti-apoptotic and Sphk2 to be pro-apoptotic^14^. Therefore, we characterized the localization of Sphk1 and Sphk2 in our pancreatic cancer cells. In HPDE, Sphk1 expression was restricted to the cytoplasm; however, all PDAC cell lines showed some degree of nuclear Sphk1 expression (Figure 2a). Interestingly, immunofluorescence of Sphk1 in BxPC-3 showed a difference in Sphk1 cellular localization dependent on the location of the cell. Cells located within the center of a growth colony showed cytoplasmic Sphk1 expression while cells at the leading edge contain nuclear Sphk1 (Figure 2b). In contrast to Sphk1, localization of Sphk2 did not change when comparing cells located within the center of a growth colony versus cells at the leading edge (Figure 2c); however, multiple PDAC cell lines showed some expression of cytoplasmic Sphk2 (Figure 2a). These results suggest that S1P signaling through S1PR1 and the translocation of Sphk1 into the nucleus and translocation of Sphk2 into the cytoplasm leads to the activation of pro-growth pathways in pancreatic ductal adenocarcinoma cells.

**Figure 2.**
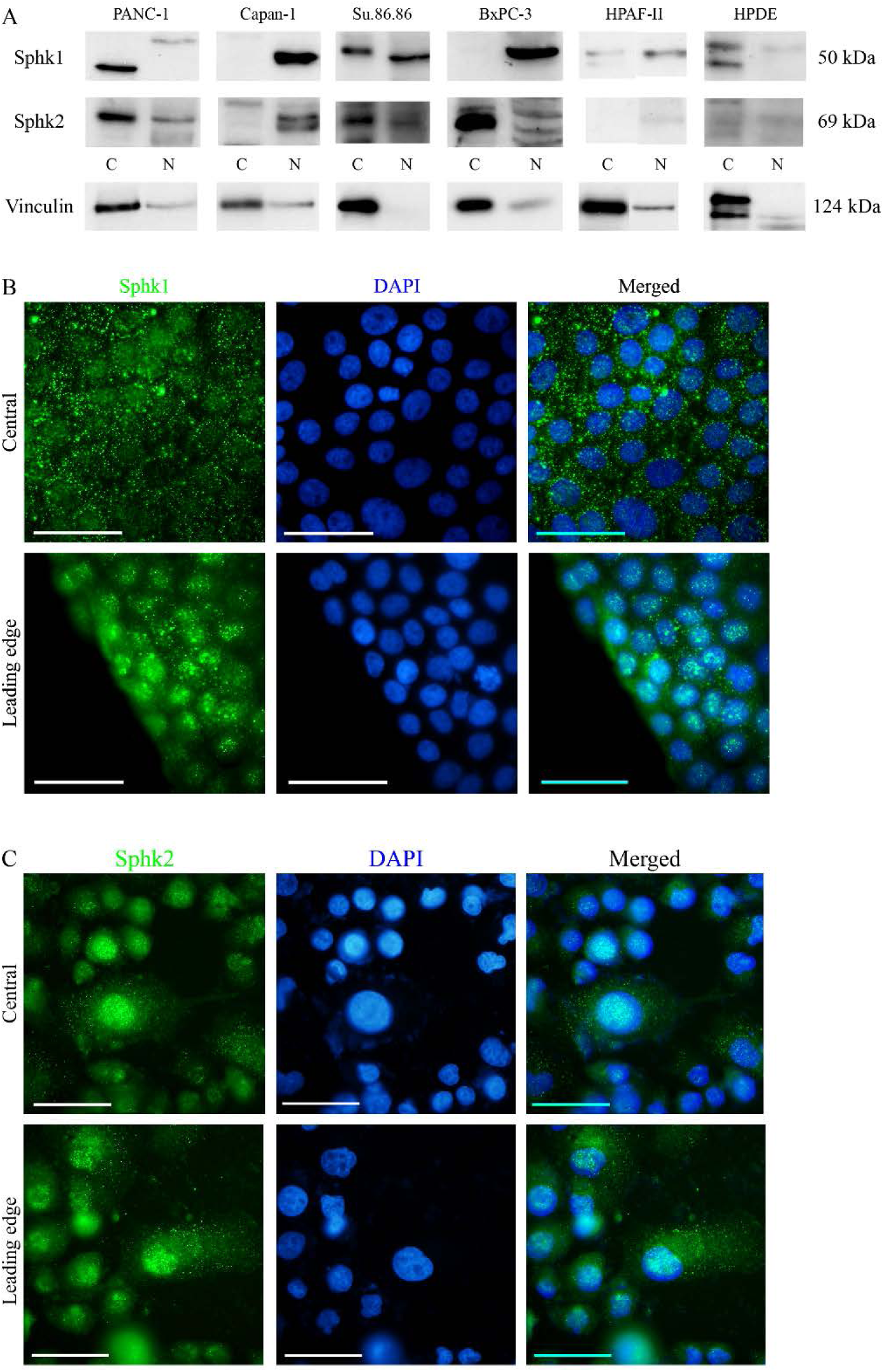
Sphk1 and Sphk2 localization is altered in pancreatic ductal adenocarcinoma. (a) Western blot analysis for Sphk1 and Sphk2. Vinculin was used as a loading control. (b) Immunofluorescence of Sphk1 in BxPC-3. (c) Immunofluorescence of Sphk2 in Bx-PC3.

### S1P modulates PDAC migration, growth, and invasion

Since S1PR1 was expressed in our panel of pancreatic ductal adenocarcinoma cell lines, we wished to determine the biological effect of S1P signaling in PDAC. BxPC-3 were grown to confluence, treated with exogenous S1P in charcoal stripped serum, and a scratch was made to measure cell growth and migration. Addition of S1P induced the cells to repopulate the wound area more rapidly than control cells in a dose-dependent manner after 24 hours (Figure 3a). This suggests that S1P induces growth of PDAC cells. Since addition of S1P led to increased migration of PDAC cells, we next determined if inhibition of S1P signaling had the opposite effect. Fingolimod (FTY720) is a functional antagonist of S1PR1 causing its degradation and has been shown to decrease proliferation and induce apoptosis in cancer cells^22-24^. To test the effect of FTY720 on cell growth, we performed an MTT assay. Treatment with FTY720 for 24 hours led to a dose-dependent inhibition of growth of PANC-1 cells (Figure 3b). Upon inspection of the cells before performing the MTT assay, there was a clear morphologic difference between PANC-1 cells treated with vehicle control versus treatment with FTY720. Treated cells had a more round shape as opposed to the more spindle-like shape of untreated cells (data not shown). Therefore, we hypothesized that inhibition of the S1P signaling pathway may alter PDAC migration and invasion. To test this, PANC-1 cells were grown in three dimensions and treated with FTY720 for 24-48 hours. After 24 hours, untreated tumorspheres grew distinctive projections into the surrounding Matrigel; however, the growth of projections was inhibited in a dose-dependent manner after treatment with FTY720 (Figure 3c). After treatment with FTY720 for 48 hours, the results were much more pronounced. These results suggest that inhibition of S1P signaling leads to a decrease in cell growth and invasion in pancreatic ductal adenocarcinoma cells potentially through modulation of the actin cytoskeleton.

**Figure 3.**
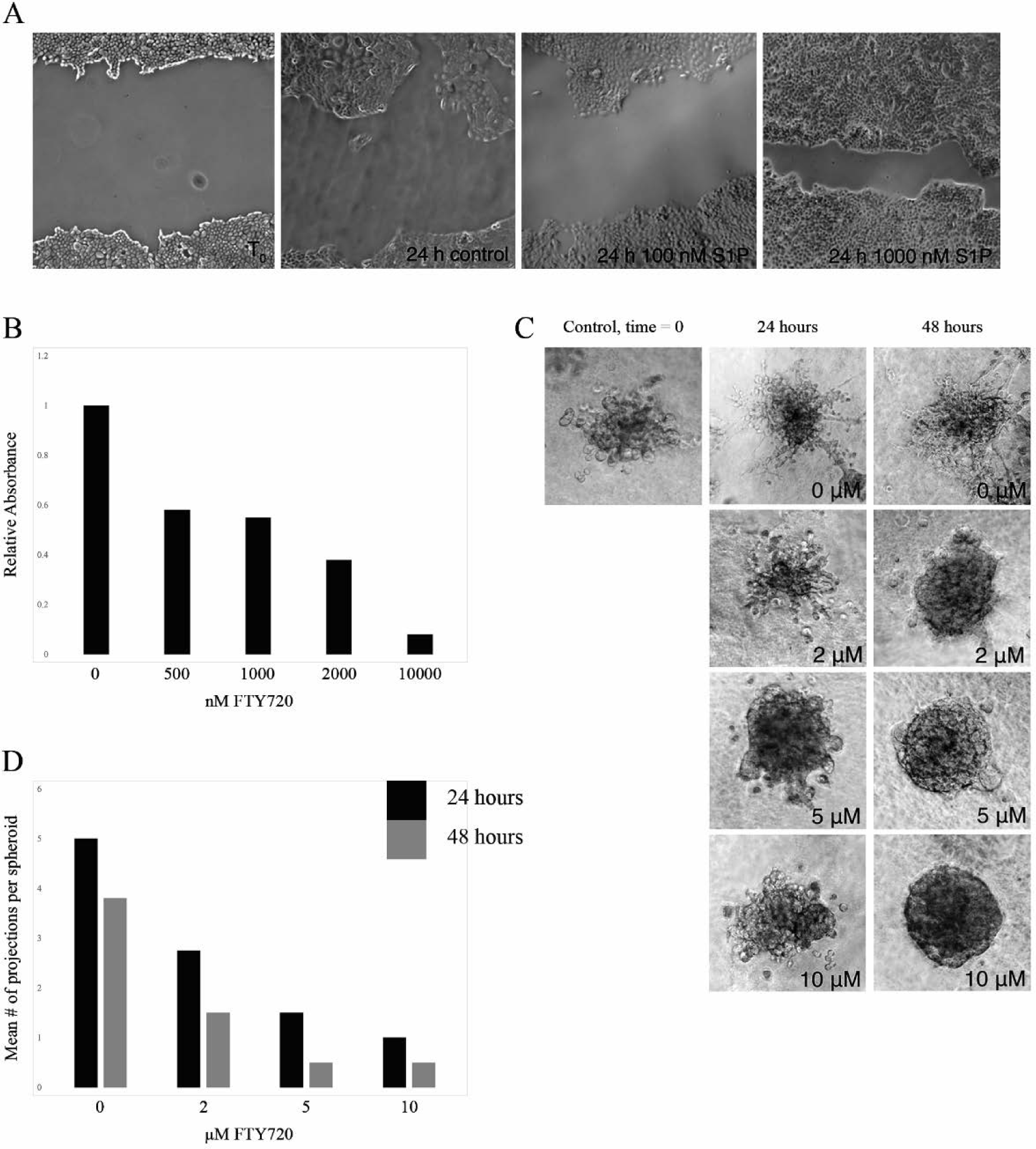
S1P modulates PDAC migration, growth, and invasion. (a) Wound healing assay using BX-PC3. Cell migration was measured using ImageJ (b) Colormetric MTT assay using PANC-1. (c) Tumorsphere assay using PANC-1. PANC-1 were suspended in 0.2% (w/v) carboxymethylcellulose and seeded at a density of 400 cells per well. Spheroid cultures were harvested and embedded into 50% Matrigel. Five separated fields were taken at 10X magnification 24 and 48 hours after embedding in Matrigel. (d) Quantification of the tumorsphere assay described in (c).

### Depletion of Sphk1 decreases PDAC cell migration and invasion

Since intracellularly generated S1P has been shown to contribute to outside-in signaling, we wanted to determine whether or not interruption of the generation of S1P leads to a similar inhibition of migration and invasion of PDAC cells. S1P destined for export from the cell is generated in the cytosol by Sphk1. Therefore, PANC-1 cells were transfected with two validated siRNAs to Sphk1 leading to an average 50% decrease in *SPHK1* mRNA expression (Figure 4a) and a > 90% decrease in Sphk1 protein expression (Figure 4b). Transfected cells were then plated in a Boyden chamber with a thin coating of Matrigel. Depletion of Sphk1 led to decreased migration and invasion of PANC-1 cells through Matrigel by approximately 50% (Figure 4c and 4d). These results suggest that members of the S1P signaling pathway may provide novel therapeutic targets to reduce PDAC cell invasion.

**Figure 4.**
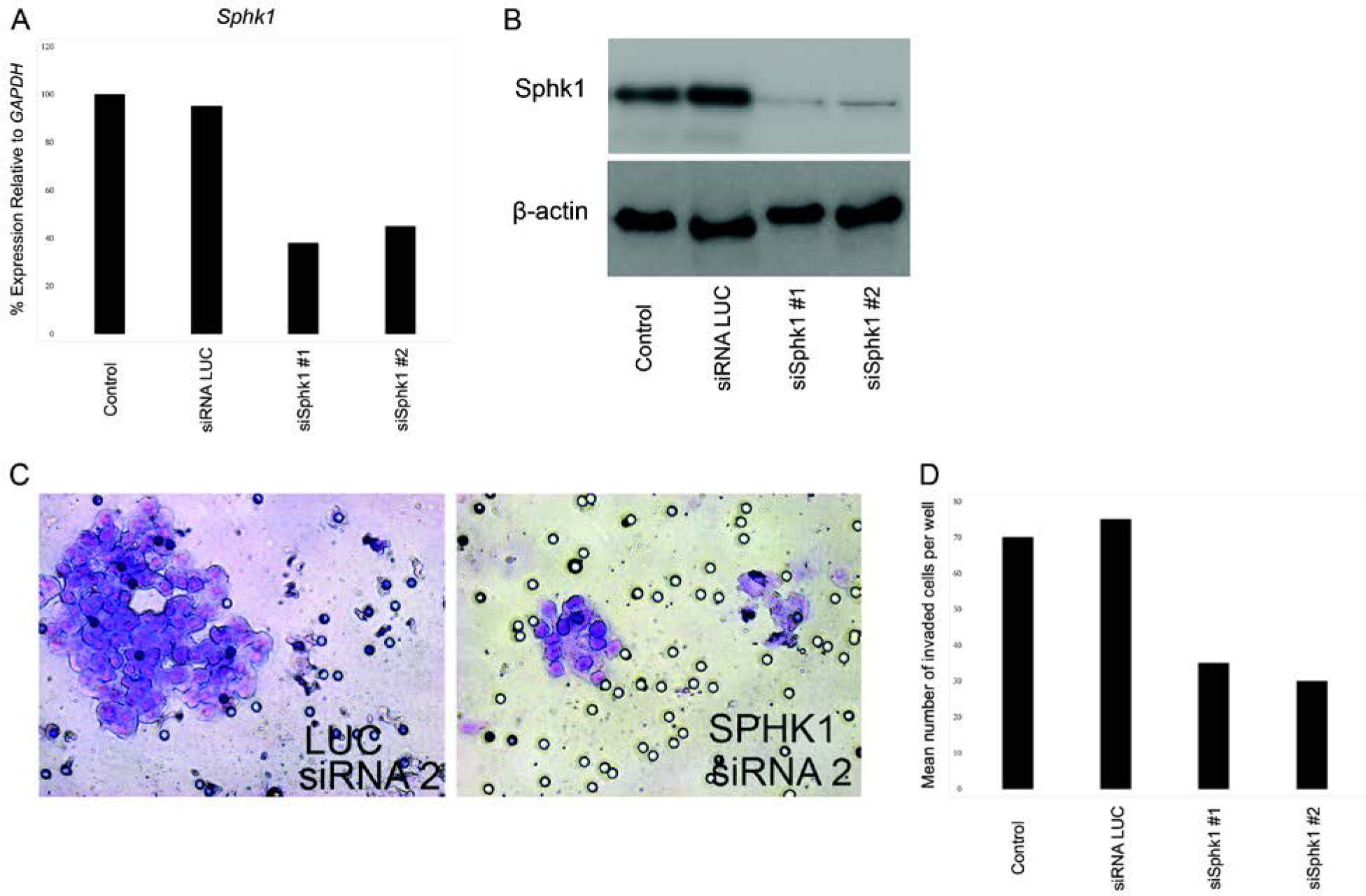
Depletion of Sphk1 decreases PDAC cell migration and invasion. (a) qPCR for Sphk1. PANC-1 cells were treated with two independent siRNAs to Sphk1 at a concentration of 50 nmol/L for 24 hours. (b) Western blot analysis for Sphk1 protein expression following siRNA transfection. (c) Boyden chamber invasion assay. siRNA transfected cells were plated in serum-free media on Matrigel-coated Boyden chambers for 24 hours and then fixed/stained with 0.5% crystal violet in 20% methanol. (d) Quantification of the invasion assay described in (c).

### S1P kinase and receptor expression in pancreatic ductal adenocarcinoma tissue

Histologic samples of patients with pancreatic ductal adenocarcinoma who underwent surgical resection were used to determine expression and localization of S1PR1, Sphk1, and Sphk2. A total of 48 different samples were used: 2 adjacent normal, 12 pancreatitis, 11 intraductal papillary mucinous neoplasm (IPMN), and 23 PDAC. As it is rare for normal pancreas to be resected, we used the samples from patients who underwent surgical resection for chronic pancreatitis as controls to calculate our composite scores. S1PR1 expression was seen in 50% of our pancreatitis samples, 80% of our IPMN samples, and 75% of our PDAC samples (Figure 5a). Although our pancreatitis samples showed the lowest percentage of S1PR1 positive cells, the composite score was not statistically significantly different than our IPMN and PDAC samples (Kruskal-Wallis, p = 0.25). Expression of Sphk1 in adjacent normal pancreas was low (Figure 5b). Additionally, there was no significant difference in the overall composite score for Sphk1 between pancreatitis, IPMN, or PDAC (Kruskal-Wallis, p = 0.35); however, localization of Sphk1 in PDAC samples was mainly cytoplasmic (Figure 5b). Like Sphk1, expression of Sphk2 in adjacent normal pancreas was low (Figure 5c). Furthermore, there was no significant difference in the overall composite score for Sphk2 between pancreatitis, IPMN, and PDAC (Kruskal-Wallis, p = 0.09); however, localization of Sphk2 in pancreatitis, IPMN, and PDAC samples was predominantly cytoplasmic as opposed to its typical location in the nucleus (Figure 5c). These results suggest that S1PR1 may play a role in the pathogenesis of PDAC while Sphk1 may be important in the progression to PDAC. The significance of altered Sphk2 localization remains unknown.

**Figure 5.**
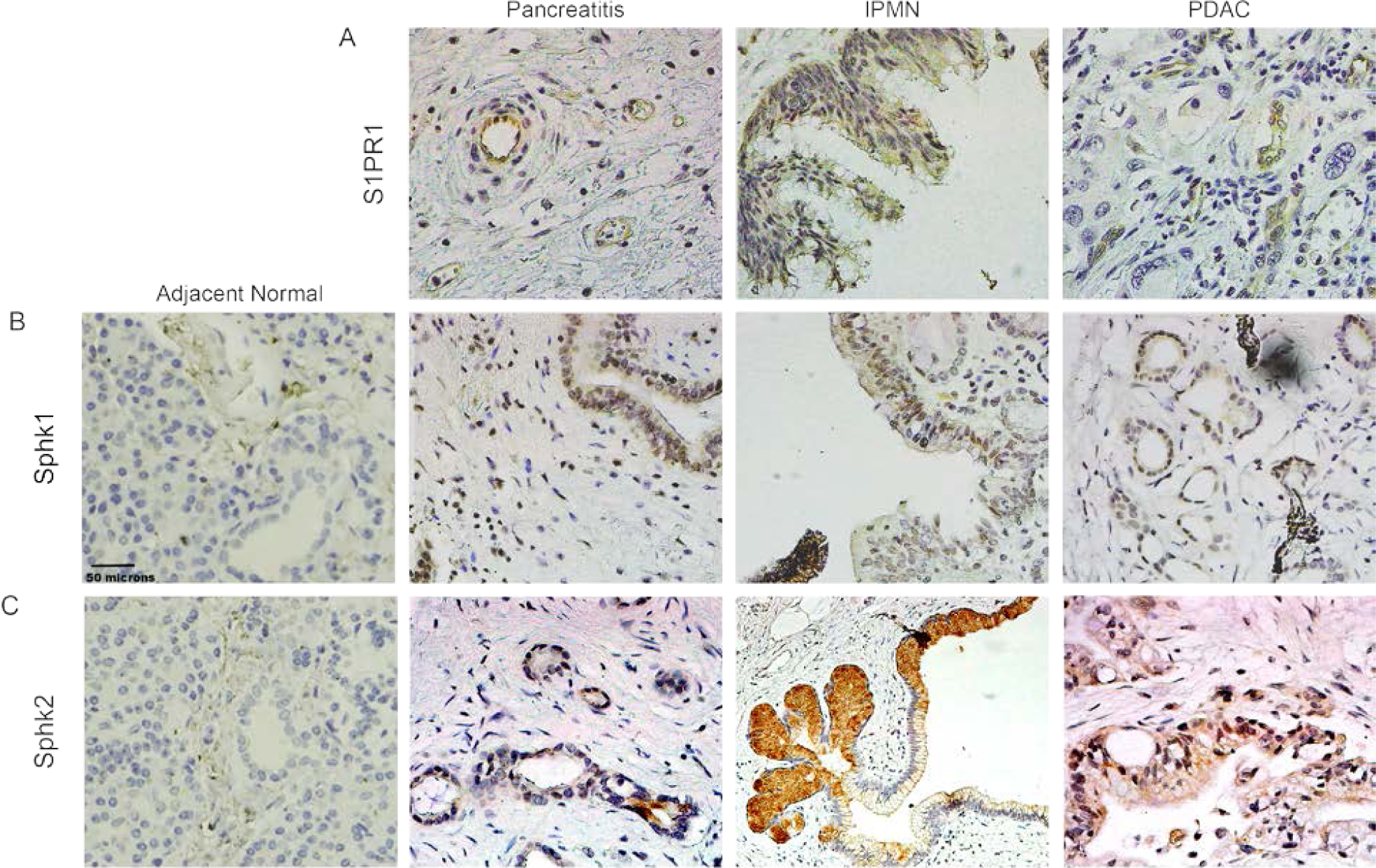
S1P kinase and receptor expression in pancreatic ductal adenocarcinoma tissue. Deidentified formalin-fixed, paraffin embedded tissue was rehydrated and stained for S1PR1, Sphk1, and Sphk2. Slides were developed with DAB and counterstained with hematoxylin. Stained slides were scanned at 40X magnification and a composite staining score was determined by multiplying the percentage of stained cells by the staining intensity. (a) IHC for S1PR1. (b) IHC for Sphk1. (c) IHC for Sphk2.

## Discussion

Sphingosine-1-phosphate has emerged as an important signaling molecule in many biological processes including growth, migration, and regulation of apoptosis; however, little is known about S1P function in the pathogenesis of pancreatic ductal adenocarcinoma. Evidence of the importance of sphingolipid metabolism has been demonstrated in the exocrine pancreas where S1P was shown to modulate inflammation in severe pancreatitis^25,26^ and gemcitabine resistance^19^. Our results demonstrate that S1PR1 is present in PDAC cell lines, but not in immortalized pancreatic ductal epithelium. Interestingly, our pancreatitis samples showed expression of S1PR1 (Figure 5); however, as PDAC arises in the context of pancreatitis, the expression of S1PR1 may be an early event in the pathogenesis of pancreatic ductal adenocarcinoma. We also noticed a trend towards further increased S1PR1 expression in PDAC samples (Figure 5) which may provide a mechanism through which pancreatic cancer becomes malignant.

Subcellular localization of sphingosine kinase is important for the phenotypic response of cells to S1P. In our study, we demonstrated altered localization of Sphk1 (Figure 2) where Sphk1 localization shifted into the nucleus in our panel of PDAC cell lines. In HPDE, bands of Sphk1 were seen in the cytoplasmic fraction suggesting cell-specific post-translational modifications that may account for altered localization. Similar results were seen for Sphk2. In normal cells, Sphk2 is typically found within the nucleus; however, cytoplasmic Sphk2 was seen in our pancreatic cancer cell lines (Figure 2). An interesting result in the present study is that the localization of Sphk1 depends on a cell’s location within a colony. Cells near the leading edge showed a clear translocation of Sphk1 to the nucleus (Figure 2) suggesting a mechanism through which altered S1P metabolism increases pancreatic ductal adenocarcinoma growth and migration.

It has been demonstrated previously that S1P promotes growth and migration in various cancers and we have shown that these events also occur pancreatic ductal adenocarcinoma. Addition of exogenous S1P led to increased migration while inhibition of S1P signaling through blockade and degradation of S1PR1 led to decreased metabolic activity and invasion (Figure 3). The most compelling results in this study are those from the three-dimensional culture experiments. Pancreatic ductal adenocarcinoma has a high propensity for invasion and metastasis due to early-onset epithelial-to-mesenchymal transition^27^. In three-dimensional culture, PDAC cells exhibit a phenotype where colonies produce protrusions from the colony that invade into the surrounding matrix; however, PDAC cells treated with fingolimod do not exhibit this invasive phenotype (Figure 3). Disruption of S1P signaling through degradation of S1PR1 showed reversal of this invasive phenotype and morphology similar to that of normal pancreatic ductal epithelium through the transition back to a more epithelial phenotype^28^ and may provide a promising therapeutic avenue for patients with metastatic disease.

Pancreatic ductal adenocarcinoma is a notoriously difficult disease to treat. Therefore, it is imperative to elucidate novel treatment modalities to improve outcomes. The results presented here offer a unique target for the treatment of PDAC. Fingolimod has already been approved by the FDA for the treatment of multiple sclerosis, is available in oral form, and has relatively few side effects. Future research including studies in animal models and clinical trials are needed to determine the efficacy of S1P inhibition as a treatment for patients with pancreatic ductal adenocarcinoma.

## Notes

### Competing Interest Statement

The authors have declared no competing interest.

